# Effects of anesthesia on ozone-induced lung and systemic inflammation

**DOI:** 10.1101/2021.12.16.472860

**Authors:** Miranda L. Wilson, Jarl A. Thysell, Kristen K. Baumann, Danny V. Quaranta, W. Sandy Liang, Michelle A. Erickson

## Abstract

Anesthetics are required for procedures that deliver drugs/biologics, infectious/inflammatory agents, and toxicants directly to the lungs. However, the possible confounding effects of anesthesia on lung inflammation and injury are underreported. Here, we tested the effects of brief isoflurane (Iso) or ketamine/xylazine/atipamezole (K/X/A) anesthesia prior to ozone exposure (4 hours, 3ppm) on lung inflammatory responses in mice. Anesthesia regimens modeled those used for non-surgical intratracheal instillations, and were administered 1-2 hours or 24 hours prior to initiating ozone exposure. We found that Iso given 1-2 hours prior to ozone inhibited inflammatory responses in the lung, and this effect was absent when Iso was given 23-24 hours prior to ozone. In contrast, K/X/A given 1-2 hours prior to ozone increased lung and systemic inflammation. Our results highlight the need to comprehensively evaluate anesthesia as an experimental variable in the assessment of lung inflammation in response to ozone and other inflammatory stimuli.

## Introduction

Ozone (O_3_) is a widespread air toxicant with well-established inflammatory effects in the lungs, but it also affects other organs, such as the liver, spleen, and brain. Approaches for selectively inhibiting lung inflammation may provide insight on the systemic effects of O_3_ and could involve direct delivery of substances to the lungs through the trachea prior to O_3_ exposure. Volatile anesthetics such as isoflurane are frequently used for intratracheal instillations and other surgeries [1–3]. However, isoflurane can have anti-inflammatory effects [4, 5], which may be most potent in the respiratory tract since it is the first site of exposure. Injectable anesthetics such as ketamine/xylazine are also used for intratracheal instillations [6], but may also modulate inflammatory responses [7–10]. Currently, little is known about how anesthesia prior to O_3_ exposure affects inflammatory responses. To test this, we exposed mice to isoflurane (Iso) or ketamine/xylazine followed by atipamezole reversal (K/X/A) prior to O_3_. Conditions of anesthesia were designed to mimic those used for non-surgical intratracheal instillations, with the rationale that pharmacological interventions to the lungs would optimally be given in a short time window prior to O_3_ exposure for maximal effects. The day after O_3_ exposure, mice were evaluated for markers of systemic and pulmonary inflammation. We found that Iso and K/X/A modify O_3_-induced lung inflammation in response to O_3_.

## Methods

### Animal Use

Female CD-1 mice, age 10-12 weeks, (Charles River Laboratories, Malvern, PA, USA) were used for air and O_3_ exposures. Females were used because they have more robust inflammatory responses to O_3_. Mice were given food and water *ad libitum* and kept on a 12 hour light/12 hour dark cycle. Protocols were approved by the institutional animal care and use committee of the VA Puget Sound Healthcare System.

### Anesthesia

Isoflurane was administered by first weighing all mice, followed by placing mice in an induction chamber filled with 4% isoflurane for 1.5 minutes, and then transferring to a nose cone where 3% isoflurane was used to maintain deep anesthesia for 5 minutes, which approximates the maximum time that a skilled technician needs to complete an intratracheal injection. Control mice were placed in the empty anesthesia chamber with room air for 1.5 minutes, and then returned to their home cage. 1-2 hours or 23-24 hours after isoflurane exposure, O_3_ exposures began. Ketamine (80mg/kg) and xylazine (10mg/kg) in sterile saline, was administered by intraperitoneal injection (IP). 20-30 minutes after the injection, mice were given 1mg/kg atipamezole IP to quickly counteract the effects of the anesthesia. Control mice were given IP saline. 1-2 hours after the atipamezole injection, O_3_ exposures began. All mice were numbered with a marker to track their identity, and mice were randomized so that the average time between anesthesia and air/ozone exposure was equivalent between groups.

### O_3_ exposure

O_3_ exposures were conducted at 3ppm for 4 hours (10:00-14:00), which induces a robust inflammatory response without inducing respiratory distress [11]. For the exposure duration, 3-4 mice were housed in standard mouse cages with free access to water, but without food or bedding to prevent consumption of ozonated food or bedding materials. Up to 4 cages at a time were placed in a 30”x 20”x 20” polypropylene chamber where O_3_ (3ppm, chamber 1) or compressed dry air (chamber 2) was pumped into the chamber at equivalent rates. Anesthesia and O_3_ exposures were replicated at least once to capture day-to-day variability, and results for each dose showed consistent trends. O_3_ levels in the chambers were generated and regulated using an Oxycycler AT42 system (BioSpherix, Parish NY, USA). Prior to each experiment, the system was calibrated using a model 106-L O_3_ detector (2B Technologies, Boulder, CO, USA), and O_3_ levels were recorded from an inlet valve in one of the O_3_ exposed mouse cages every 10 seconds for the duration of exposures. In all experiments, O_3_ achieved its target concentration within 10 minutes, and levels were regulated within 10% of the target concentration (3ppm +/− 0.3ppm) thereafter.

### Blood Collection and Serum Amyloid A measurement

22-24 hours after the start of O_3_ exposure (12:00-14:00), mice were re-weighed and deeply anesthetized with IP urethane (Millipore Sigma, St. Louis, MO, USA). Blood was collected from the abdominal aorta, allowed to clot for 30 minutes, and then placed on ice. Blood was centrifuged at 2500g for 15 minutes and serum was collected, aliquoted, and frozen at −80°C. Serum was diluted 1/10,000 for mice exposed to ozone and 1/50 for mice not exposed to ozone. SAA mouse duoset kits (R and D systems, Minneapolis, MN, USA) were used to quantify serum amyloid A (SAA) in serum.

### BAL and pulmonary inflammation assessment

Bronchoalveolar lavage (BAL) was performed on mice by perfusing and aspirating the lungs through the trachea 3 times with 1 ml sterile phosphate buffered saline (3 mL total). BAL fluid was stored on ice and centrifuged at 200g for 5 minutes at 4°C. The supernatant was removed and 0.5 mls of supernatant was reserved to resuspend the cell pellet. The remainder was frozen for measurement of total BAL protein, which was performed using a microBradford assay. Total cells were manually counted using a hemacytometer. Differential cell counts were performed on Hemacolor-stained cytocentrifuge preparations. Cell counts were performed using Image J, and at least 200 cells were counted to determine relative cell proportions.

### Statistics

Statistical analysis was done with Prism 8.4.2 (Graph-Pad Software, San Diego, CA, USA). Data are reported as mean ± SD and analyzed by two-way ANOVA for main effects and interactions and Tukey’s multiple comparisons test for comparisons of group means.

## Results

### Effects of isoflurane (Iso) and ketamine/xylazine/atipamezole (K/X/A) anesthesia on ozone-induced pulmonary inflammation and injury

We first determined whether O_3_-induced responses in the lungs differed with Iso or K/X/A. Since drugs or other interventions would ideally be given a short time before exposure to ozone to maximize their effects, we compared Iso and ketamine at 1-2 hours prior to ozone exposure, and also tested a 24 hour Iso pre-exposure to determine whether effects were sustained (Schematics in Figure 1A and B). Although the controls for ketamine were IP injected with saline, whereas Iso controls were not, we found that IP injections did not significantly influence the measured outcomes, and so we combined the Iso and Ketamine no anesthesia control groups for analysis. O_3_ significantly increased BAL total cells, macrophages, and neutrophils except in the 1-2 hour Iso group (Figure 2A-C). We further show that mice exposed to K/X/A and O_3_ had significantly increased BAL total cells, macrophages, and neutrophils vs. other O_3_-exposed groups. O_3_ significantly increased BAL protein in the no anesthesia group and 23-24h Iso groups, but not in the 1-2 hour Iso or K/X/A groups (Figure 2D). The F-statistics and p-values for main effects and interactions are reported in Table 1.

**Figure 1.**
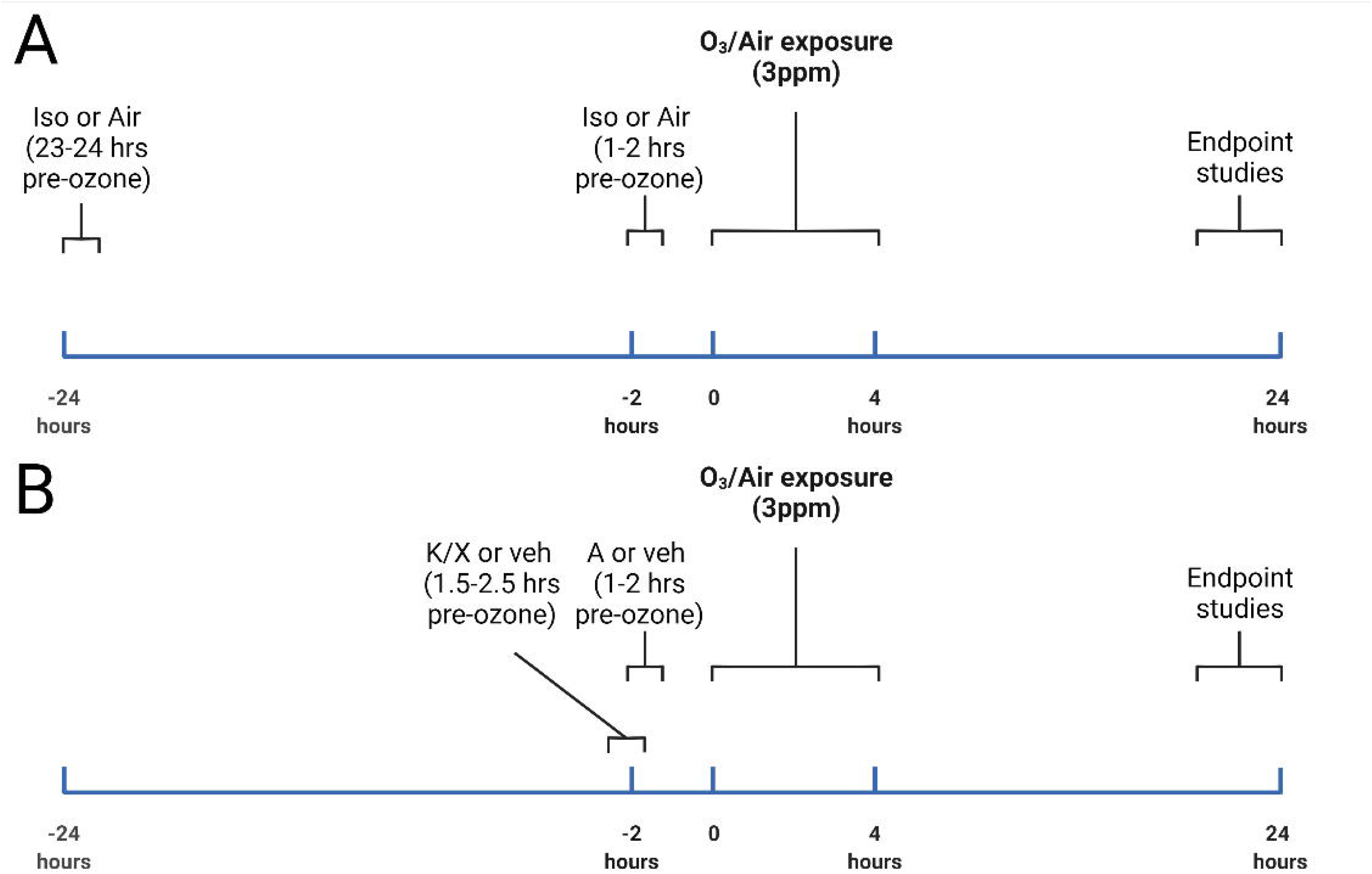
Schematic of anesthesia/ozone (O_3_) exposure protocols. Figure 1A shows the exposure protocol for mice pre-exposed to isoflurane (Iso) or air control, and Figure 1B shows the exposure protocol for mice pre-exposed to Ketamine/Xylazine (K/X) and atipamezole (A) or saline vehicle. Created with Biorender.com.

**Figure 2.**
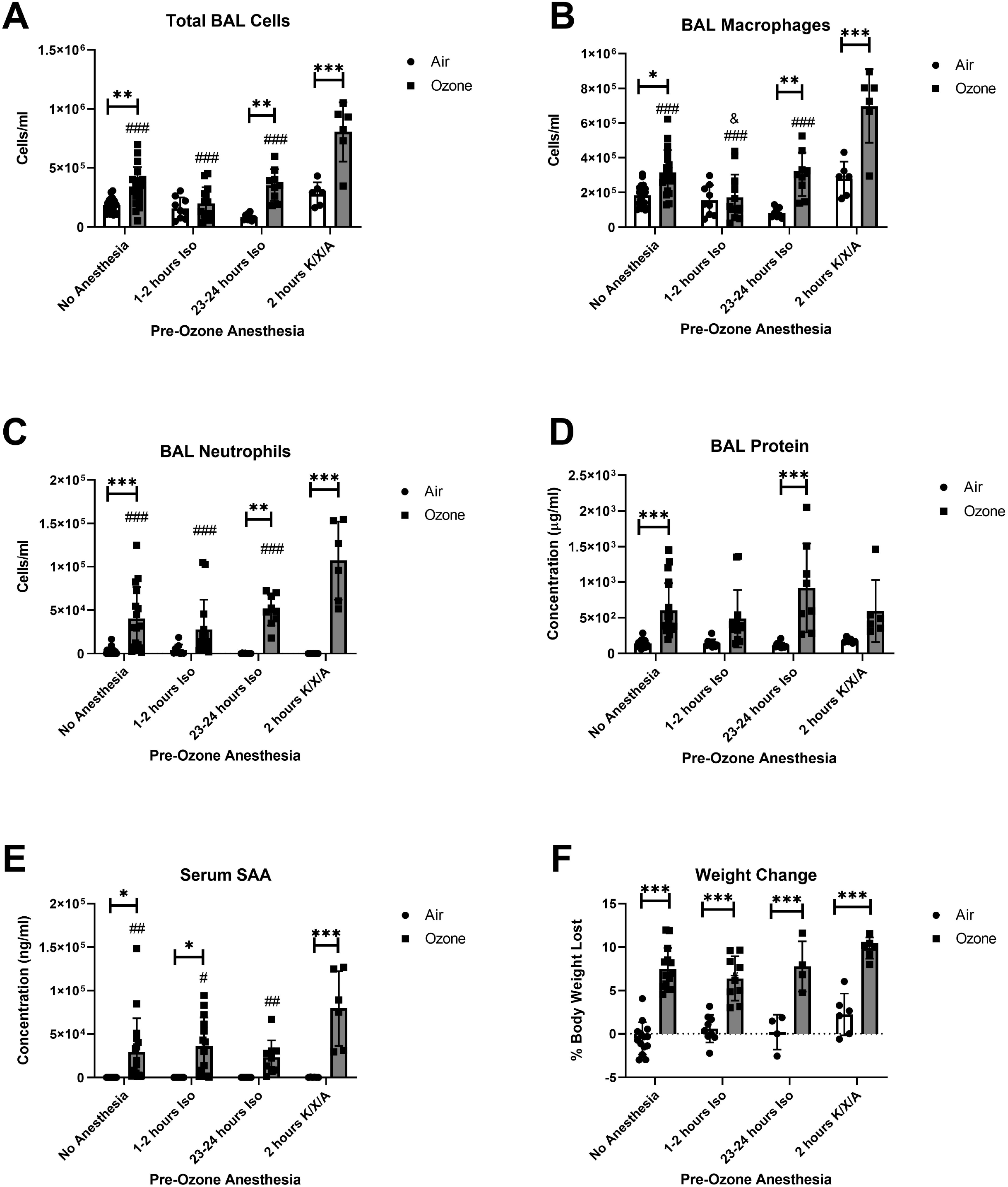
Effects of Isoflurane (Iso) and ketamine/xylazine/atipamezole (K/X/A) on inflammatory (A-C) and vascular damage (D) markers in the lung, blood levels of serum amyloid A (SAA, E), and weight loss (F). In all panels, n=6-20 per group. ***p < 0.001, **p < 0.01, *p < 0.05, air vs. O_3_; ###p < 0.001, ##p < 0.01, #p < 0.05, vs. K/X/A + O_3_ group. &p < 0.05 vs. No Anesthesia + O_3_. Group mean comparisons were carried out using Tukey’s multiple comparisons test.

**Table 1.** Two-Way ANOVA analysis results showing percentages of variation by factor (main effects, interactions), and F-statistics. For Figure 2D, one sample was excluded because of apparent blood contamination. For Figure 2E, one sample was excluded because not enough serum was recovered to complete the assay. For Figure 2F, body weights were not measured in one experimental replicate and so 23 samples are missing.

| Figure | Anesthesia | Exposure | Interaction |
| --- | --- | --- | --- |
| 2A - Total<br>BAL Cells | 28.97%; F (3,82) =<br>22.15, p < 0.0001 | 29.96%; F (1,82) =<br>68.73, p < 0.0001 | 12.79%; F (3,82) =<br>9.780, p < 0.0001 |
| 2B - BAL<br>Macrophages | 31.77%; F (3,82) =<br>23.49, p < 0.0001 | 25.59%; F (1,82) =<br>56.74, p < 0.0001 | 11.72%; F (3,82) =<br>8.662, p < 0.0001 |
| 2C - BAL<br>Neutrophils | 9.766%; F (3,82) =<br>6.535, p = 0.0005 | 44.88%; F (1,82) =<br>90.08, p < 0.0001 | 11.82%; F (3,82) =<br>7.907, p = 0.0001 |
| 2D - BAL<br>total protein | 2.776%; F (3, 81) =<br>1.309, p = 0.2773 | 33.06%; F (1, 81) =<br>46.76, p < 0.0001 | 3.888%; F (3,81) =<br>1.833, p = 0.1478 |
| 2E - Serum<br>SAA | 7.296%; F (3,81) =<br>3.668, p = 0.0156 | 35.35%; F (1, 81) =<br>53.31, p < 0.0001 | 7.204%; F (3,81) =<br>3.622, p = 0.0165 |
| 2F - Weight<br>change | 7.784%; F (3, 58) =<br>6.412, p = 0.0008 | 52.74%; F (1, 58) =<br>130.3, p < 0.0001 | 1.903%; F (3,58) =<br>1.567, p = 0.2070 |

### Effects of Iso and K/X/A anesthesia on ozone-induced systemic inflammation and weight loss

We next determined whether the effects of anesthesia extended to markers of ozone-induced systemic inflammation, and behavioral effects. We have previously shown that the acute phase protein serum amyloid A (SAA) is consistently increased in blood following a 3ppm exposure to O_3_, and its upregulation occurs in the absence of changes in other pro-inflammatory cytokines in blood [12]. Weight loss is a measurable outcome of sickness behavior, which can be induced by O_3_. Anesthesia has no effect on O_3_-induced weight loss based on the post-hoc mean comparisons of the ozone groups (Figure 2E), however there was a significant main effect of treatment that explained 5.172% of the total variation (Table 1). O_3_ significantly increased SAA levels in all groups except for mice exposed to Iso 23-24 hours prior (Figure 2F). SAA levels induced by O_3_ were significantly higher in mice pre-anesthetized with K/X/A vs no anesthesia or Iso.

## Discussion

We found that isoflurane inhibits O_3_-induced pulmonary inflammation if given 1-2 hours prior to exposure, and this effect diminishes by 24 hours. Isoflurane induces anesthesia via potentiation of inhibitory γ-aminobutyric acid type A (GABA A) receptors and inhibition of glutamatergic α-amino-3-hydroxy-5-methyl-4-isoxazolepropionic acid (AMPA) and N-methyl-d-aspartate (NMDA) receptors [13]. Isoflurane also has antiinflammatory activities in a variety of rodent models [4], although the mechanisms are not well-defined. It has been shown that activation of GABA A receptors and inhibition of NMDA receptors have anti-inflammatory effects on immune cells [14–16], suggesting that the anti-inflammatory activities of Iso could be mediated through direct actions on leukocytes. In contrast to Iso, K/X/A increased O_3_-induced inflammatory responses to O_3_. Anti-inflammatory/protective effects through NMDA receptor antagonism have been described for ketamine and ketamine/xylazine combinations [9]. Whereas xylazine is an α2 adrenergic receptor agonist and potentiates anesthesia, atipamezole is an antagonist, and leads to rapid anesthesia reversal within about 20 minutes [17]. The data on pro-vs. anti-inflammatory activities of the α2 adrenergic receptor are equivocal, however a recent study showed that atipamezole may promote or prolong inflammation through its action via an α2-independent mechanism [18]. Future work is needed to delineate the immunomodulatory mechanisms of Iso and K/X/A.

Our results are the first to show that isoflurane exposure prior to O_3_ inhibits pulmonary inflammation, whereas K/X/A exposure has an enhancing effect. These results complement other recent works that have investigated anesthesia’s effects on pulmonary inflammation in response to ozone and other pro-inflammatory agents [5, 16, 19]. Although the scope of our study is limited in that we did not define a full time course of anesthesia’s effects, our results highlight that brief, acute, anesthetic regimens can alter inflammatory responses to O_3_. We conclude that anesthetics are important experimental variables whose evaluation is necessary for studies requiring their use prior to exposure to O_3_ or other pathological stimuli.

## References

1. K. E. Driscoll, D. L. Costa, G. Hatch, R. Henderson, G. Oberdorster, H. Salem, et al. Intratracheal instillation as an exposure technique for the evaluation of respiratory tract toxicity: uses and limitations. Toxicol Sci. 2000;55(1):24–35.

2. L. Morales-Nebreda, M. Chi, E. Lecuona, N. S. Chandel, L. A. Dada, K. Ridge, et al. Intratracheal administration of influenza virus is superior to intranasal administration as a model of acute lung injury. J Virol Methods. 2014;209(116–20.

3. M. B. Lawrenz, R. A. Fodah, M. G. Gutierrez, and J. Warawa. Intubation-mediated intratracheal (IMIT) instillation: a noninvasive, lung-specific delivery system. J Vis Exp. 2014;93):e52261.

4. Y. M. Lee, B. C. Song, and K. J. Yeum. Impact of Volatile Anesthetics on Oxidative Stress and Inflammation. Biomed Res Int. 2015;2015(242709.

5. S. E. Lacher, C. Johnson, F. Jessop, A. Holian, and C. T. Migliaccio. Murine pulmonary inflammation model: a comparative study of anesthesia and instillation methods. Inhal Toxicol. 2010;22(1):77–83.

6. M. N. Helms, E. Torres-Gonzalez, P. Goodson, and M. Rojas. Direct tracheal instillation of solutes into mouse lung. J Vis Exp. 2010;42).

7. S. Tan, Y. Wang, K. Chen, Z. Long, and J. Zou. Ketamine Alleviates Depressive-Like Behaviors via Down-Regulating Inflammatory Cytokines Induced by Chronic Restraint Stress in Mice. Biol Pharm Bull. 2017;40(8):1260–1267.

8. M. H. Chen, C. T. Li, W. C. Lin, C. J. Hong, P. C. Tu, Y. M. Bai, et al. Rapid inflammation modulation and antidepressant efficacy of a low-dose ketamine infusion in treatment-resistant depression: A randomized, double-blind control study. Psychiatry Res. 2018;269(207–211.

9. M. K. Erdem, G. Yurdakan, and E. Yilmaz-Sipahi. The effects of ketamine, midazolam and ketamine/xylazine on acute lung injury induced by alphanaphthylthiourea in rats. Adv Clin Exp Med. 2014;23(3):343–51.

10. J. Szelenyi, J. P. Kiss, E. Puskas, Z. Selmeczy, M. Szelenyi, and E. S. Vizi. Opposite role of alpha2- and beta-adrenoceptors in the modulation of interleukin-10 production in endotoxaemic mice. Neuroreport. 2000;11(16):3565–8.

11. S. Kierstein, K. Krytska, S. Sharma, Y. Amrani, M. Salmon, R. A. Panettieri, Jr., et al. Ozone inhalation induces exacerbation of eosinophilic airway inflammation and hyperresponsiveness in allergen-sensitized mice. Allergy. 2008;63(4):438–46.

12. M. A. Erickson, J. Jude, H. Zhao, E. M. Rhea, T. S. Salameh, W. Jester, et al. Serum amyloid A: an ozone-induced circulating factor with potentially important functions in the lung-brain axis. FASEB J. 2017;31(9):3950–3965.

13. N. J. Michelson and T. D. Y. Kozai. Isoflurane and ketamine differentially influence spontaneous and evoked laminar electrophysiology in mouse V1. J Neurophysiol. 2018;120(5):2232–2245.

14. M. G. Reyes-Garcia, F. Hernandez-Hernandez, B. Hernandez-Tellez, and F. Garcia-Tamayo. GABA (A) receptor subunits RNA expression in mice peritoneal macrophages modulate their IL-6/IL-12 production. J Neuroimmunol. 2007;188(1-2):64–8.

15. A. K. Bhandage, Z. Jin, S. V. Korol, Q. Shen, Y. Pei, Q. Deng, et al. GABA Regulates Release of Inflammatory Cytokines From Peripheral Blood Mononuclear Cells and CD4(+) T Cells and Is Immunosuppressive in Type 1 Diabetes. EBioMedicine. 2018;30(283–294.

16. Y. Wang, S. Yue, Z. Luo, C. Cao, X. Yu, Z. Liao, et al. N-methyl-D-aspartate receptor activation mediates lung fibroblast proliferation and differentiation in hyperoxia-induced chronic lung disease in newborn rats. Respir Res. 2016;17(1):136.

17. L. Mees, J. Fidler, M. Kreuzer, J. Fu, M. T. Pardue, and P. S. Garcia. Faster emergence behavior from ketamine/xylazine anesthesia with atipamezole versus yohimbine. PLoS One. 2018;13(10):e0199087.

18. J. Hu, S. Vacas, X. Feng, D. Lutrin, Y. Uchida, I. K. Lai, et al. Dexmedetomidine Prevents Cognitive Decline by Enhancing Resolution of High Mobility Group Box 1 Protein-induced Inflammation through a Vagomimetic Action in Mice. Anesthesiology. 2018;128(5):921–931.

19. K. Yuki, Y. Mitsui, M. Shibamura-Fujiogi, L. Hou, K. C. Odegard, S. G. Soriano, et al. Anesthetics isoflurane and sevoflurane attenuate flagellin-mediated inflammation in the lung. Biochem Biophys Res Commun. 2021;557(254–260.

